# Three-Dimensional Impedance Tomographic Mapping of Metabolically Active Endolumen

**DOI:** 10.1101/2020.09.24.312025

**Authors:** Parinaz Abiri, Yuan Luo, Zi-Yu Huang, Mehrdad Roustaei, Sandra Duarte-Vogel, Quinyu Cui, René R. Sevag Packard, Ramin Ebrahimi, Peyman Benharash, Yu-Chong Tai, Tzung K. Hsiai

**Author notes:** These authors contributed equally to this work (Parinaz Abiri, Yuan Luo, Zi-Yu Huang).

## Abstract

Real-time detection of vulnerable atherosclerotic lesions, characterized by a high content of oxidized low-density lipoprotein (oxLDL)-laden macrophages or foam cells, remains an unmet clinical need. While fractional flow reserve (FFR)-guided revascularization in angiographically intermediate stenoses is utilized to assess hemodynamic significance, *in vivo* detection of oxLDL-rich plaques may provide a new paradigm for treating metabolically unstable lesions. Herein, we have demonstrated endoluminal mapping of lipid-laden lesions using 3-D electrical impedance spectroscopy-derived impedance tomography (EIT) in a pre-clinical swine model. We performed surgical banding of the right carotid arteries of Yucatan mini-pigs, followed by 16 weeks of high-fat diet, to promote the development of lipid-rich lesions. We implemented an intravascular sensor combining an FFR pressure transducer with a 6-point micro-electrode array for electrical impedance spectroscopy (EIS) measurements. 3-D EIT mapping was achieved using an EIS-based reconstruction algorithm. We demonstrated that EIT mapping corresponds to endoluminal histology for oxLDL-laden lesions. We further used computational models to theoretically predict and validate EIS measurements. Thus, our 3-D EIS-derived EIT provides *in vivo* detection of metabolically active plaques with the goal of guiding optimal intravascular intervention.

**One Sentence Summary:** This work demonstrates *in vivo* mapping of oxidized LDL-laden endolumen by deploying an intravascular dual-sensor to a swine model of atherosclerosis.

## Introduction

Cardiometabolic syndromes, including hyperlipidemia, obesity, and diabetes, constitute a rising epidemic in the United States. These often silent disorders are associated with chronic diseases, including atherosclerosis (*1*). A subgroup of atherosclerotic lesions are known to spontaneously rupture, leading to myocardial infarction and stroke (*2, 3*); however, reliable detection of vulnerable plaques is yet to be realized clinically.

Metabolically active plaques consist of a thin fibrous cap, oxidized lipids, and M1 macrophages (*4-11*). Plaque rupture occurs when the fibrous cap overlying the lipid-laden lesion is biomechanically disrupted in the presence of shear stress, thus exposing the thrombogenic subendothelial factors to the bloodstream, resulting in platelet adhesion, activation, and aggregation (*12*). Various catheter-based techniques, including intravascular ultrasound and near infrared spectroscopy, have been developed for the characterization of arterial plaques. Measurement of Fraction Flow Reserve (FFR) (*13*), defined as the ratio of pressure across the stenotic lesions (P_downstream_/P_upstream_) during coronary catheterization (*14-16*), is often employed to assess hemodynamically significant lesions deemed to be of intermediate severity (*17-19*). However, the predictors for metabolically active, albeit non-obstructive, lesions prone to rupture remain undefined by FFR, resulting in a false negative rate of over 20% (*20*).

We have previously established the sensitivity and specificity of electrical impedance spectroscopy (EIS) for the detection of oxidized low-density lipoprotein (oxLDL)-laden macrophages in a rabbit model of atherosclerosis (*21-23*). This method was demonstrated by integrating three intravascular sensing modalities; namely, shear stress sensors, intravascular ultrasound (IVUS), and EIS (*24-28*). This integration enabled sequential detection of disturbed blood flow, plaque visualization by IVUS, and oxLDL-laden lesions by EIS (*21, 28-30*). Oxidized lipid in macrophages has been shown to destabilize the fibrous cap by activation of matrix metalloproteinases. These oxLDL-rich arterial walls exhibit a significant increase in the frequency-dependent impedance magnitude by EIS interrogation (*22, 23, 31*).

In this context, we sought to demonstrate 3-D impedance mapping of oxidized lipid-laden carotid arteries in a pre-clinical model using our 3-D EIS-derived impedance tomography (EIT). Lipid-rich plaques were created in the Yucatan mini-pigs via surgical banding of right carotid arteries to induce disturbed flow, followed by 16-weeks of high-fat diet to promote the development of atherosclerosis. Next, we deployed the dual FFR-EIS sensor, including a pressure transducer and 6-point microelectrodes, to interrogate changes in endoluminal impedance in the right and left (sham) carotid arteries. We reconstructed the 3-D impedance mapping derived from EIS measurements, demonstrating correlation with prominent E06 staining for oxLDL-rich lesions in the right carotid arteries. We further simulated the EIS measurements by using 3-D histology-derived finite element models with the assigned electrical properties to collagen, lipid, and smooth muscle. This computational model allowed for theoretical prediction and validation of the EIS measurements. Our results demonstrated that EIS-based *in vivo* detection of lipid-rich endolumen may represent a new paradigm for identifying metabolically active, albeit angiographically non-obstructive lesions.

## Results

### A. Swine Model for OxLDL-Rich Atherosclerotic Lesions

Six Yucatan mini-pigs were fed a high-fat diet for 16 weeks, followed by an interrogation of the right common carotid artery. All animals were monitored over the 16-week course via CT imaging for the progression of carotid stenoses. Baseline (0 weeks), intermediate (8 weeks), and terminal (16 weeks) diameter measurements of the stenosed right carotid artery demonstrated atherosclerotic formation as compared to the left carotid artery (**Figure 1**). The average internal diameters in the stenosed right carotids were reduced by ∼33% (from 4.5 mm to 3 mm), whereas the diameter of left (control) carotid arteries increased from 4.6 mm to 5.0 mm, likely due to a compensatory response to the decreased flow in the right.

**Fig. 1.**
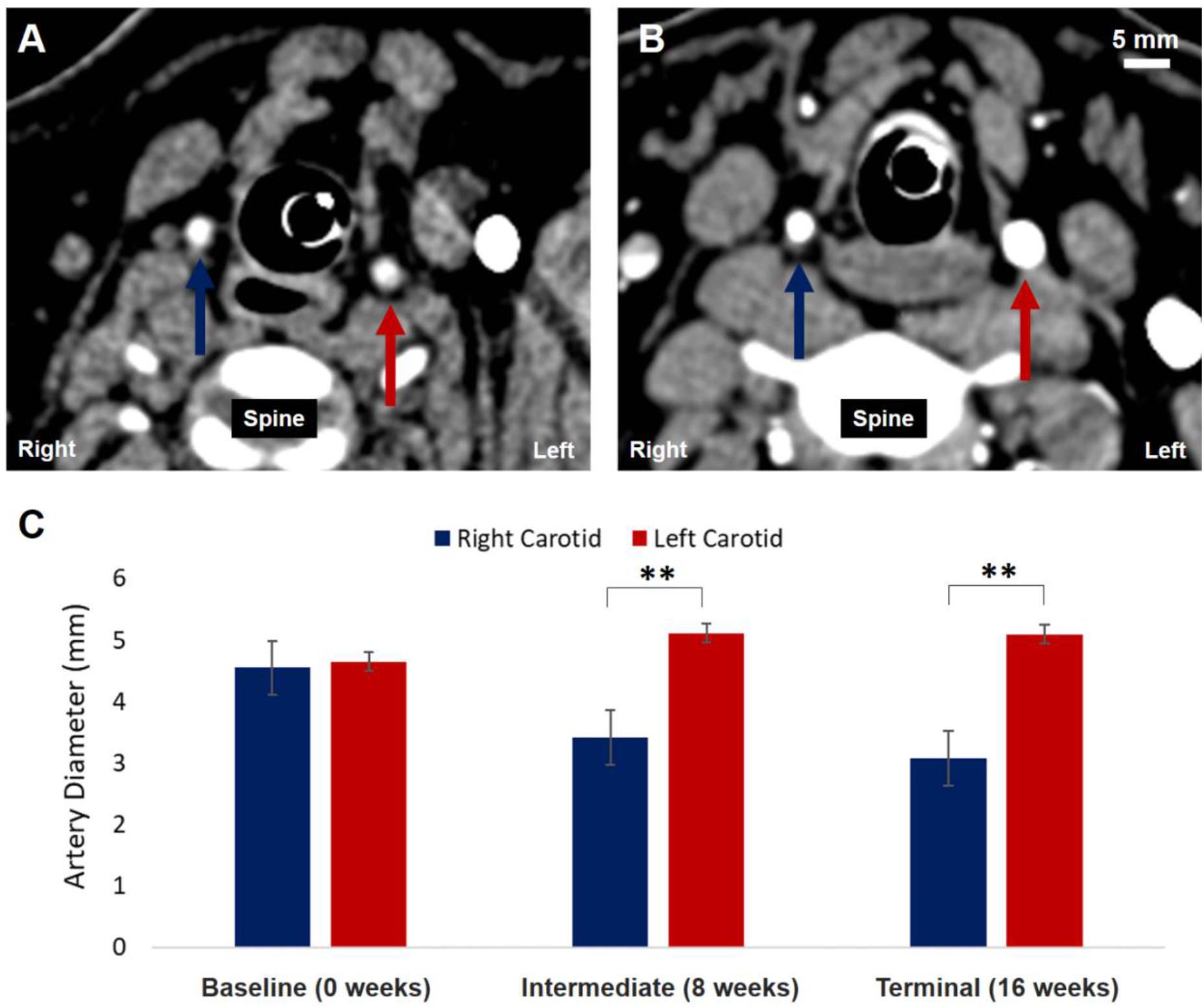
Carotid Artery Stenosis. (A) Baseline axial CT of carotid arteries (CA) demonstrate a similar diameter between the right and left CA. (B) Terminal axial CT demonstrates a decrease in the right CA diameter as compared to the left. (C) Comparison of the mean carotid artery diameters at 0 weeks (baseline), 8 weeks post-surgery (intermediate), and 16 weeks post-surgery (terminal) demonstrates no significant difference at baseline (*p* > 0.05, n=6), but significant difference at intermediate and terminal time (*p* < 0.05, n=6).

### B. Correlation between EIS-derived EIT mappings and 3-D Histology in the oxLDL-Laden atherosclerotic lesions

The dual EIS-FFR sensor was deployed to the carotid arteries of the Yucatan mini-pigs (**Figure 2**). To demonstrate the pressure measurements for FFR, we recorded pressure fluctuation starting from the sensor insertion up to the stenotic lesion (**Supplementary SI-1**). Using the 6-electrode array, we performed EIS measurements between any pair of electrodes, with 15 permutations, including 3 permutations to link the vertically aligned electrodes, 6 permutations to link the circumferentially paired, and 6 permutations to link the diagonally paired (**Supplementary Figure SI-5A)**.

**Fig. 2.**
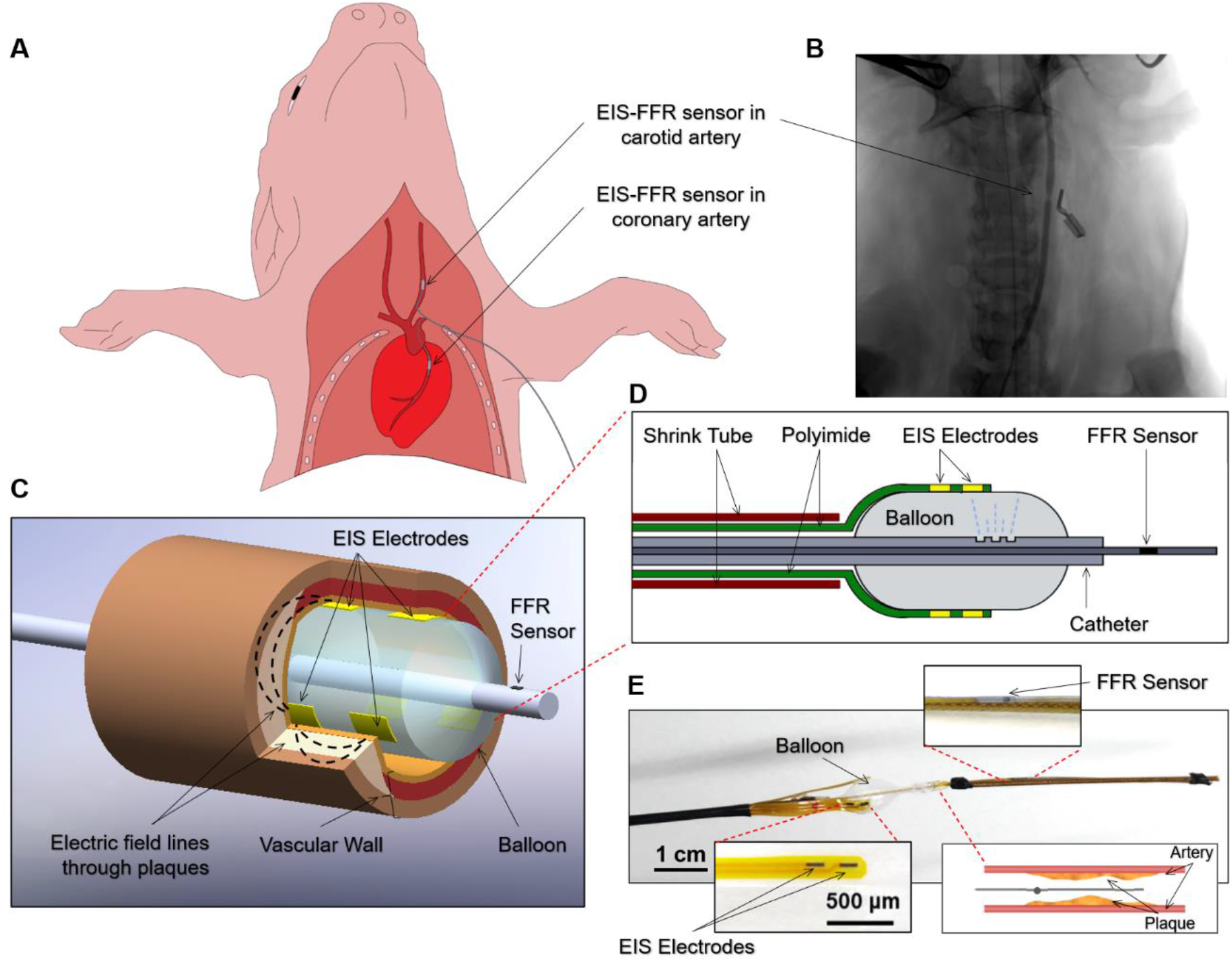
Device Schematic. (A) Schematic illustrates the right carotid arteries for the deployment of 6-point electrodes for EIS and FFR. The dual EIS-FFR sensor is designed for intravascular deployment to either the carotid or coronary arteries in the Yucatan min-pigs. (B) Following surgical banding of the right carotid artery and 16 weeks of high-fat diet, the radiographic image demonstrates the deployment of the dual EIS-FFR sensor to the stenotic lesions (lateral radiopaque marker identifies the position of the lesions). (C) Schematic of dual EIS-FFR sensor illustrates the interrogation of an atherosclerotic lesion. The flexible 6-point EIS electrodes were fixated on the balloon, generating 15 pairs of EIS measurements. An FFR pressure sensor was upstream from the balloon. (D) Cross-sectional view shows the inflated balloon. The flexible polyimide EIS electrodes were fixated on the balloon, and the FFR pressure sensor was cannulated through to the catheter. Micro holes were opened on the catheter. (E) A photo of the dual-sensor catheter provides the position of the EIS electrodes fixated to the balloon in relation to the FFR pressure sensor as well as the catheter in relation to the plaque.

We measured baseline EIS profiles in the carotid arteries prior to balloon inflation (**Figure 3A, black series**). The EIS impedance profiles of the right carotid artery (with stenosis) were consistently higher than those of left carotid artery (control) following balloon inflation (**Figure 3A, colored series**). Next, we reconstructed the EIS-derived EIT mappings with the 15 impedance values at 10 kHz for the individual carotid arteries (**Figure 3B**). We observed a correlation between the EIT mapping and 3-D histology, as supported by the Movat stain for tissue composition, E06 for oxLDL, and 3-D histology reconstruction (**Figure 3C-E**). The EIT mapping of the left carotid artery shows a yellow to orange color-coded gradient, indicating the absence of oxLDL (**Figure 3, left column**). In the right carotid #1 (RC1), the dark brown color-coded gradients align with the prominent semi-circumferential E06 staining (**Figure 3, middle column**). In the right carotid #2 (RC2), the dark brown gradients correspond to the presence of E06 staining in the right upper quadrant of the carotid circumference (**Figure 3, right column**). The prominent oxLDL distribution in RC1 further corroborated a broader range of impedance profiles than that of RC2.

**Fig. 3.**
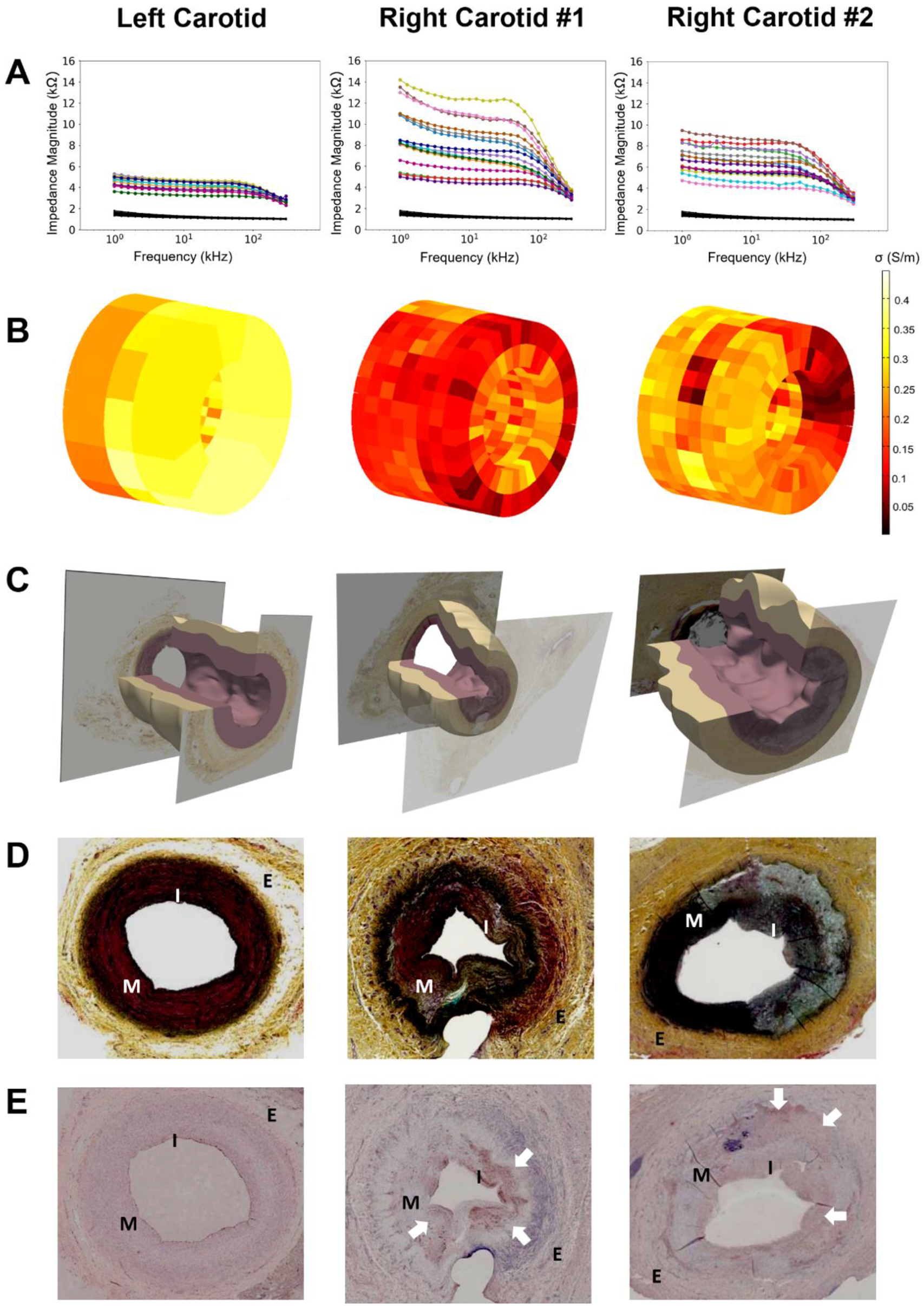
EIS-derived 3-D EIT Mapping of Carotid Artery. Data from a representative left (sham) and two right carotid arteries. (A) Frequency-dependent EIS profiles from 1 to 300 kHz were compared between the left and right carotid arteries. Baseline EIS profiles (black) were acquired prior to balloon inflation. The individual EIS profiles reflect 15 permutations from the 6-point electrodes. A total of 26 impedance measurements from 1 to 300 kHz were obtained for each EIS profile. (B) 3-D EIS-derived EIT of the tunica intima and tunica media were constructed from the impedance profiles at 10 kHz. (C) The 3-D histological reconstruction recapitulates the endoluminal topology from 11 cross-sections of a segment (4 mm) of carotid arteries. (D) The representative Movat staining for connective tissue was compared between the left and right carotid arteries. (E) The representative E06 staining for oxidized LDL was also compared. The white arrows point to the presence of oxLDL. I: tunica intima; M: tunica media; E: tunica externa.

### C. Computational Modeling to Validate Experimental EIS Profiles

Histological data and 3-D computational modeling were utilized to validate our experimental impedance data. Through this effort, we identified the electrode positions (z, θ) in relation to the lumen by comparing the computational data with experimental EIS profiles (**Figure 4A**). Next, we compared the measured impedance values with computational outputs at 10 kHz; namely, left carotid (LC), RC1, RC2 at Position A (RC2-A), and RC2 at Position B (RC2-B) (**Figure 4B**).

**Fig. 4.**
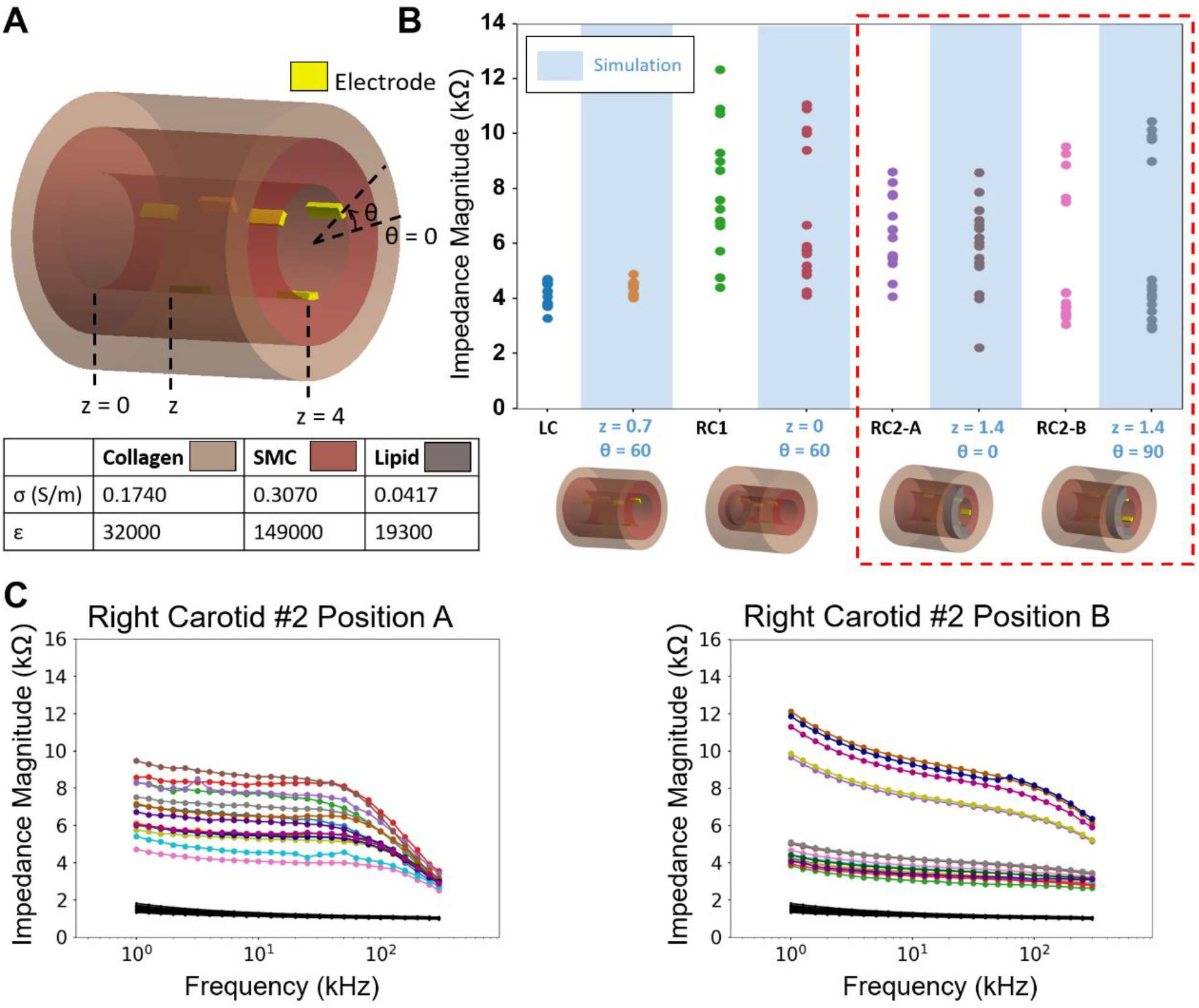
Comparing the Computational with Measured EIS Profiles. (A) The position of the electrodes in the 3-D model is defined by the polar coordinates, where *z* is defined as the distance between the edge of the electrodes and *z* = 0, and *θ* is the angle subscribed between the midpoint of the first pair of the electrodes and the *θ* = 0. (B) The measured impedance values at 10 kHz were compared with the computational (simulation) impedance values. The 3-D schematics of the arterial segments illustrate the position of the electrodes in the lumen. (C) The 15 impedance profiles were compared between two different rotational positions of the electrodes from the same lumen.

The computational values were in agreement with the low impedance values in LC as compared to both RC1 and RC2. We further demonstrated that changes in the electrode positions and rotations imparted no significant shift to the EIS profiles in LC. The model also predicted the distributions of impedance measurements in RC1 and RC2. The different electrode positions in relation to the oxLDL-laden lumen engendered a distinct distribution of EIS profiles between RC2-A and RC2-B (**Figure 4C**). In the presence of different electrode positions to the lipid-laden lumens, RC2-B data revealed two distinct regions with broader impedance spectra as compared to RC2-A data. By simulating these changes, we identified two combinations of *z* and *θ* values for reconstructing the impedance distributions that overlapped with the EIS measurements from RC2-A and RC2-B, respectively. Computational impedance profiles from additional combinations of electrode position are illustrated in **Supplementary SI-2**.

## Discussion

Our investigation on EIS-derived EIT mapping introduces a transition from 2-D intravascular spectroscopy to 3-D tomography. This pre-clinical model of atherosclerosis recapitulated the oxidized lipid-laden endolumen for EIT mapping in the Yucatan mini-pigs. The 3-D histology-derived computational model simulated impedance distributions to predict and validate the measured EIS profiles. The dual EIS-FFR sensor, integrated with 6-electrode and the pressure transducer, further facilitated concurrent measurements while reducing radiation exposure and procedural duration. Overall, EIT mapping for *in vivo* detection of lipid-rich endolumen may represent a new strategy to identify metabolically active, albeit angiographically non-obstructive lesions.

Numerous techniques have previously been developed to assess plaque vulnerability. The standard-of-care technique of angiography, now combined with fractional-flow reserve (FFR) measurements, provides valuable information regarding the hemodynamic significance of lesions. However, FFR alone is unable to detect plaque composition or vulnerability to spontaneous rupture. Non-invasive imaging modalities, including Computed tomography (CT) and magnetic resonance imaging (MRI), identify stenotic lesions, but with inadequate spatial resolution and specificity. To acquire high resolution images, Li *et al* developed an integrated IVUS and optical coherence tomographic (OCT) catheter (*32*). While the combined technique allows for improved plaque characterization, OCT is limited by the need for saline solution flushing (*32*). Photoacoustics takes a different approach to plaque analysis by imaging the vasa vasorum and intraplaque micro-vessels using high photo-absorption and thermal expansion of blood (*33-37*). However, the heat generated from thermal expansion may have adverse effects on the vulnerable plaque and saline flushing is indicated. Near-infrared fluorescence (NIRF) has been used as an indicator of inflammation (*38*), and [^18^F]-fluorodeoxyglucose (^18^FDG) is capable of demonstrating metabolic activity by positron emission tomography (PET) (*39*). However, the injection of contrast agents is required for NIRF and injection of radioactive isotopes for PET imaging. Alternatively, microbubble contrast agents are utilized in acoustic angiography to assess lesion microvasculature (*40-43*); however, the need for cessation of blood flow during this procedure significantly limits its clinical application and safety. In this context, the integration of FFR with 3-D EIS mapping may provide an efficient strategy to characterize the metabolically active lesions during diagnostic angiography.

As a catheter-based approach, EIS has the capability to identify oxLDL-laden macrophages in the subendothelial layers of atherosclerotic lesions. To this end, we demonstrated the capability of 3-D EIS-derived EIT for detecting the gradients of lipid-laden plaques in the mini-pig model of carotid atherosclerosis. The deployment of the dual EIS-FFR sensor enabled our fundamental EIS measurements to characterize fatty streaks in the *ex vivo* human arch and the eccentric atherosclerotic lesions in the New Zealand White rabbit model (*21-23, 31*). In this study, the carotid arteries of the mini-pigs were selected for comparable diameters (3-5 mm) to human coronary arteries (*44*). The surgical banding of right carotid arteries, followed by diet-induced hyperlipidemia, accelerated the development of metabolically active plaques (*45*). Thus, we were able to demonstrate the pre-clinical feasibility of EIT to interrogate the atherosclerotic lesions in the swine model.

Notably, our EIS-derived EIT algorithm directly solved the forward problem by obviating the inverse ill-posed problem encountered by other EIT algorithms (*46-48*). We used the “genetic algorithm” to optimize the conductivity distributions that were closely overlapping with the Movat staining for connective tissue and E06 staining for oxLDL, as represented by the color-coded gradients of 3-D mapping (**Figure 3**) (*49, 50*). For these reasons, EIT facilitates the characterization of the oxidative state (oxidized LDL) of the metabolically active lesions during a diagnostic angiogram.

Our histological segmentation and 3-D reconstruction of the carotid arteries also allowed for the creation of a computational model to validate the measured EIS profiles. Electrical impedance values are governed by the distinct tissue composition and precise boundary conditions of the organ system (*22*), and insufficient knowledge of arterial wall composition and topography may deviate the computational modeling from the experimental measurement (*51*). For this reason, we obtained multi-slice and axial histology to establish a 3-D arterial computational model with well-defined layers of tissue properties (*σ* = tissue conductivity and *ε* = permittivity), including collagen, fatty tissue, and smooth muscle (**Supplementary Table I**). In the computational model, we simulated the changes in the electrode positions in relation to the arterial wall. Based on observations from the multi-slice histology, the semi-circular region of the arterial wall with the prominent oxLDL staining (**Figure 3D & E**) likely contributed to the increase in impedance. By comparing the position of the electrodes in the cylindrical coordinates (*z, θ*), we identified two distinct electrode positions to reconstruct a comparable impedance distribution from two oxLDL-laden carotid arteries (**Figure 4B**). Hence, the computational models simulated two different electrode positions to predict and validate the 3-D EIS measurements.

Both EIS-derived EIT and histology-derived computational model provide complementary and synergistic insights into the endoluminal metabolic state. However, effective reconstruction of the 3-D EIS-derived EIT is dependent on the number of electrodes. The current 6-electrode configuration may be expanded to 12 electrodes to enhance the spatial resolution. While increasing the number of electrodes would require additional computation for tomographic reconstruction, our current methodology provides a foundation for this future step. Furthermore, our computational model has provided validation of our experimental EIS measurements. With the *a priori* knowledge of the boundary condition and histology, the computational model simulated the changes in electrode positions from two separate EIS measurements; thus, supporting the current 6-point electrode configuration to map the oxLDL-rich endolumen.

In sum, we have demonstrated the deployment and implementation of a single intravascular dual-sensor that integrates a pressure sensor with a multi-electrode configuration for 3-D EIS measurements. Our acquired electrical impedance tomography of the artery exhibits a high correlation with the 3-D histology in the pre-clinical model of atherosclerosis, supporting the detection of oxidized LDL-laden plaques with high-risk features. Future studies aimed at refining this integrated technique may greatly advance our clinical understanding of the vulnerable plaque and have implications in interventions aimed at plaque modification.

## Materials and Methods

### A. Device Micro-Fabrication and Integration

A catheter-based dual EIS-FFR sensor (7F diameter) was developed for intravascular delivery (Figure 2). Custom-made flexible polyimide electrodes (600 µm x 300 µm) were designed and manufactured (FPCexpress, Canada). Six of these electrodes were positioned in two rows (hence 3 by 3 electrodes) at 1.4 mm apart along the circumference of an inflatable balloon (very-low-durometer urethane Ventiona Medical, NH) whose length is 9 mm, and the inflatable diameters range from 2 to 10 mm (Figure 1C-E). These two rows of electrodes were fixated onto the balloon using the silicone adhesive. The balloon was coaxially inserted into the distal end of a polyethylene catheter (Vention Medical, NH), and was anchored with the epoxy glue. Micro holes were opened on the catheter to allow for balloon inflation. A pair of tantalum foils (Advanced Research Materials, UK) was added to both ends of the balloon as radiopaque markers. The catheters were insulated with heat-shrink tubing (Vention Medical, NH) (Figure 2D). The electrical conduction to the impedance analyzer was connected by soldering a joint between the copper wires (26 AWG) and exposed contact pads at the terminal end of the flexible electrodes. A commercial fractional flow reserve (FFR) probe (St. Jude Medical, MN) was coaxially inserted into the catheter and hermetically sealed to the ends of the catheter with epoxy. The electrodes were electroplated with platinum black (Sigma-Aldrich) to increase the junction capacitance and to enhance the accuracy of two-point electrode measurements.

### B. Measurement System Design

Alternating Current (AC) with peak-to-peak voltages of 50 mV and sweeping frequencies ranging from 1 – 300 kHz were applied to acquire the impedance measurements (Gamry Series G 300 potentiostat, USA). We acquired ten impedance values per frequency decade. A manual syringe inflator with a pressure gauge was used to ensure reproducible balloon inflation for EIS measurements of the endolumen.

Pressure measurements for FFR were acquired by adapting the commercial FFR sensor through a custom-built Wheatstone bridge (see **Supplementary SI-3**). Individual resistors were chosen according to the intrinsic resistance of the pressure sensor components; a linear relationship between voltage change and pressure difference was confirmed. The entire set-up for pressure measurements was calibrated via a commercial pressure sensor (LPS331AP, STMicroelectronics, Switzerland).

### C. A Swine Model of Atherosclerosis

A combination of high-fat diet and carotid arterial banding was previously demonstrated to promote the initiation of atherosclerosis in a swine model (46). After surgically-induced stenosis and diet-induced hyperlipidemia, plaques preferentially develop in the vessel wall proximal to the stenoses where disturbed flow or oscillatory shear stress developed (46, 52). For this reason, we compared the right (banding) with left (sham) carotid arteries using 3-D EIS-derived EIT mapping in the Yucatan miniature pigs (n = 6, 20-30 kg; S & S Farms, Ranchita, CA). The animal study was approved by the UCLA Office of Animal Research in compliance with the institutional IACUC protocols. The surgical procedures and the postoperative care were performed by experienced veterinarians from the Division of Laboratory Animal Medicine at UCLA School of Medicine.

All animals were fed on a high-fat diet containing 4% cholesterol, 20% saturated fat, and 1.5% supplemental choline (Test Diet; Purina, St. Louis, MO) for 2 weeks before surgical banding of the right carotid arteries. The pigs were anesthetized with intramuscular Tiletamine and Zolazepam, and Isoflurane was given to maintain general anesthesia during the procedure. A 6F introducer sheath was inserted percutaneously via the Seldinger procedure into the right or left femoral artery to monitor blood pressure and to provide access for angiography. Bupivacaine was subcutaneously injected in the ventral neck along the path of the incision site. A midline skin incision was placed at the neck. Both right and left common carotid arteries were dissected approximately 5 cm in length, but the right common carotid artery was tied off with a suture (Ethicon, Cornelia, Ga) around a spacer (approximately 1.3 mm in diameter) positioned on the external surface of the artery. The spacer was subsequently pulled out, leaving a 50-70% stenosis (**Figure 5A**). Postoperative CT angiography was performed to monitor the degree of surgical stenosis (**Figure 5B**).

**Fig. 5.**
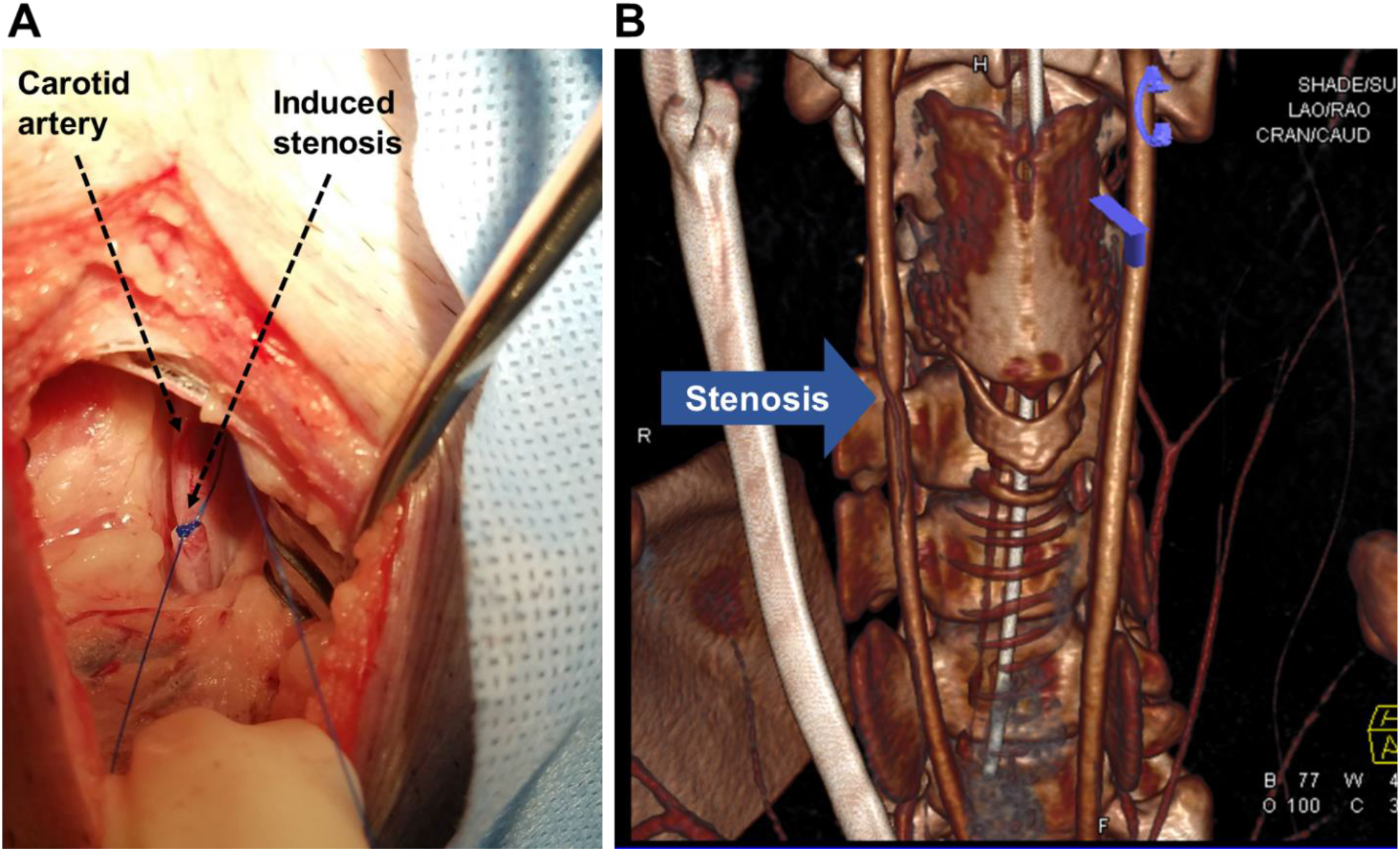
Surgical Banding of the Right Carotid Artery. (A) A midline skin incision was made at the neck. The right common carotid artery was dissected from the tissue, and tied off to create a 50-70% stenosis in the mid-segment of the artery. (B) 3-D CT angiogram reveals the stenotic right common carotid artery in comparison to the left.

A sham operation was performed on the left carotid artery and control by dissecting through a midline skin incision at the neck under the general anesthesia. The surgical wound was closed layer by layer to avoid manipulation to the adjacent tissues. The animals were allowed to recover after surgery, and they were resumed to the high-fat diet for 16 weeks. Serial aortic CT angiograms were performed to assess the diameters of the carotid arteries following Iodinated contrast injection to the tail vein at baseline, 8 weeks, and 16 weeks.

### D. Deployment of dual EIS-FFR Catheter to interrogate lipid-rich arterial wall

To deploy the dual EIS-FFR sensor for interrogation of right carotid arteries, the animals were anesthetized as described above. Bupivacaine was subcutaneously injected in the ventral neck along the path of the incision site. A midline skin incision was placed at the neck. The common carotid arteries were dissected and a surgical cut-down was performed to directly introduce the sheath and device into the carotid artery at the site of stenosis in the right carotid artery and at the approximate mirror location in the left carotid artery. For the right carotid artery, the EIS-FFR sensor was advanced to the level of the stenosis with the radiopaque marker under fluoroscopic guidance (Siemens Artis Zeego with robotic arm) (**Figure 2E**), and iodinated contrast dye was injected into the vessel.

The pressure reading from the FFR was recorded from the insertion of the sensor until it was positioned near the lesions. Next, the balloon with the six circumferentially fixed electrodes was inflated to a constant pressure at ∼14 psi via a mechanical pump to be in contact with the endoluminal surface. EIS measurements were conducted using the Gamry system in which fifteen scans for each EIS sensor were performed based on 15 paired combinations between the six electrodes. A peak-to-peak voltage of 50 mV was delivered to acquire the frequency-dependent impedance profiles ranging from 1 – 300 kHz. We acquired 10 data points per frequency decade. Following the interrogation of lipid-laden and control arteries, the catheter was removed, and the pigs were euthanized with an overdose of pentobarbital and phenytoin. Bilateral carotid arteries were collected for histology and immunohistochemistry.

During the sensor deployment, two of the six mini-pigs developed device-related carotid embolization, thus preventing EIS measurement and histological analysis. In another animal, electrode contamination distorted the data collection. One of the six animals did not develop grossly visible plaque in the right carotid artery. For these reasons, we ultimately collected EIS measurements from two animals with angiographically visible lesions in the right carotid arteries with the corresponding control carotid artery.

### E. 3-D EIS-derived EIT Mappings from 15 EIS Impedance Profiles

3-D EIS-derived EIT mappings allowed for visualization of the endoluminal conductivity or impedance distributions in terms of EIS impedance profiles. Impedance computation was performed at 10 kHz based on the fading of the electrode contact impedance beyond 1 kHz. This frequency reflected the resistance contribution from the collagen, lipid, and smooth muscle in the arterial wall (**Supplementary SI-4**). The reactance contribution was considered to be negligible. To reconstruct EIS-derived EIT mappings, we divided the arterial segments into 864 elements, of which the smooth muscle cell layer was represented by 576 elements (**Supplementary Figure SI-5B**). Assuming each element to be uniform, we derived the initial impedance/conductivity for the smooth muscle from the EIS measurements. We used the conductivity of collagen for the remainder of the elements, and we computed the impedance values for the 15 permutations via the EIDORS (version 3.8) (53). We incorporated this information and the “genetic algorithm” to alter the conductivity value of each element, and we created a new set of impedance values for the 15 permutations (49, 50) (refer to **Supplemental SI-5**). The conductivity maps were generated by minimizing the impedance difference between the measured and computed data. The details for deriving the conductivity values were provided in **Supplementary SI-5**.

### F. Histology, Immunostaining, and Reconstruction of 3-D Histology

The carotid arteries with stenosis were dissected into segments at 10 mm in length. The samples were prepared in 10% formalin, embedded in paraffin, and sectioned from the center with 5 slices on each side at 0.4 mm apart. A total of 11 slices were sectioned, and each slice was further sectioned into thin sections at 5 µm in thickness for (1) Movat staining for the connective tissue, including elastic fibers (black), collagen and reticular fibers (yellow), fibrin (bright red), and muscle (red); and (2) E06 staining for oxidized-LDL-laden lesions (dark red). The immunohistochemistry was performed by the CV Path Institute, Inc. (Gaithersburg, MD, USA).

In addition, we reconstructed the 3-D histology from these 11 slices to model the complete segment of the dissected carotid artery. The histological slices were aligned using the center of the cross-sectional images, and were inputted into Image-J software (National Institute of Health, Bethesda, MD, USA). Segmentation of the lumen, media, and adventitia layer was performed in SimVascular and the gaps between the slices were interpolated using the spline function (54). The results were exported to Paraview for 3-D visualization (55).

### G. 3-D histology-derived EIS Profiles to Predict and Validate EIS measurements

While 3-D mapping identifies the lipid-rich lesions, the number of electrodes influences the spatial resolution of the impedance tomography. We validated the 3-D EIS profiles via finite element simulation with the aforementioned 3-D histology. To simulate the endoluminal topology, we utilized multi-slice histological sections as established by the 3-D model in Comsol Multiphysics.

As illustrated by the Movat staining, individual slices from the carotid artery were divided into 3 layers; namely, lumen, inner, and outer arterial wall (**Supplementary SI-6)**. Our immunohistochemistry (**Figure 3D-E**) revealed that the outer wall was comprised of mostly collagen (yellow), the inner wall was mostly smooth muscle cells (red), and a segment of the inner wall was prominent for oxLDL (white). The 2-D outline of each layer was first extruded from the histological slices in AutoCAD, and was stacked to reconstruct a 3-D model with the center in alignment with the geometric center of the lumen. While the lumen was deformed by the inflated balloon (∼1 cm long) during the EIS measurement, we approximated a uniform circle for all cross-sections. The lumen circumference was estimated from the average circumference of each slice (**Supplementary SI-6**).

In the absence of any a priori knowledge of the precise position of the electrodes relative to each artery, we scanned a wide range of electrode positions to optimally reproduce the measured EIS values. The electrode positions in the cylindrical coordinates corresponded to the distance between the edge of the electrodes and z = 0, z, and to the rotational angle, θ. We used 3 different z values (0, 0.7, and 1.4) and 4 different θ values (0°, 30°, 60°, and 90°) to generate 12 possible electrode positions for each arterial model.

The computational EIS model was governed by the Time-Harmonic Maxwell equation. Assuming a negligible contribution from the magnetic field (46), we arrived at the following expression:

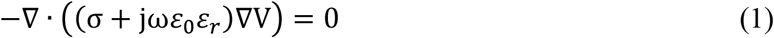

For each combination of *z* and *θ*, 15 impedance values (from the permutations of the 6 electrodes) were solved numerically by using the Comsol AC/DC module in conjunction with the assigned material properties; namely, conductivity (*σ*) and permittivity (*ε*) for the collagen, lipid, and smooth muscle (**Supplementary SI-6, Table I**).

We reconstructed the individual arterial models from the histology, and we compared between experimental and computational EIS profiles to identify the probable electrode position during the experiments. We adopted the following mathematical criteria for the identification of the closest alignment: the impedance values from the experiments were sorted in the ascending order: *Z*_*exp*,1_, *Z*_*exp*,2_, *Z*_*exp*,3_, … …, *Z*_*exp*,15_. For each combination of the electrode positions, the impedance values were sorted analogously: *Z*_*sim*,1_, *Z*_*sim*,2_, *Z*_*sim*,3_, … …, *Z*_*sim*,15_. Next, we compared the summation of the square of the differences between the experimental and simulated EIS as follows:

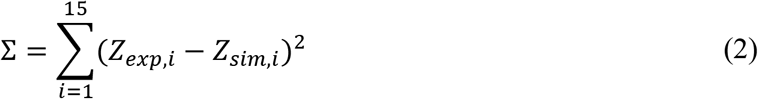

The combination of electrode placement resulting in the smallest summation was taken to be the best fit of the measured impedance values.

## Acknowledgments

We appreciate Chadi Nahal for sorting the histological data.

## Author contributions

PA, YL, and ZYH designed and performed the experiments, and they wrote the manuscript. PA also contributed to data integration and revision. YL also fabricated the device and performed the data analysis. ZYH further performed the computational modeling. MR contributed to the 3-D histology for modeling and simulation of deployment. SDV helped with the planning and deployment of sensors to the pre-clinical model. QC helped with the illustrations. RRSP helped with the planning of pre-clinical studies, imaging, and connecting with CV path for histology. RE and PB helped with the clinical correlation and manuscript revision. YCT supervised the microfabrication of the catheter-based sensors and data analyses. TKH conceived, implemented, and supported the project, and he revised the manuscript.

