## Supplementary Information for "Three-Dimensional Impedance Tomographic Mapping of Metabolically Active Endolumen"

Manuscript Supplementary

**Supplementary Materials:**

*SI-1. Pressure Monitoring*

As a demonstration of the pressure recording function of the dual-function device, we recorded the changes in pressure before the device was inserted, and we continued after the device was positioned at the lesions (**Fig. SI-1)**. An average increase in pressure of 40 mmHg was recorded from both the right and left carotid arteries. The low-pressure value was consistent with the partial blockage of blood flow in the presence of a balloon catheter, which was positioned upstream from the pressure sensor of the FFR probe (see **Figure 2E**).

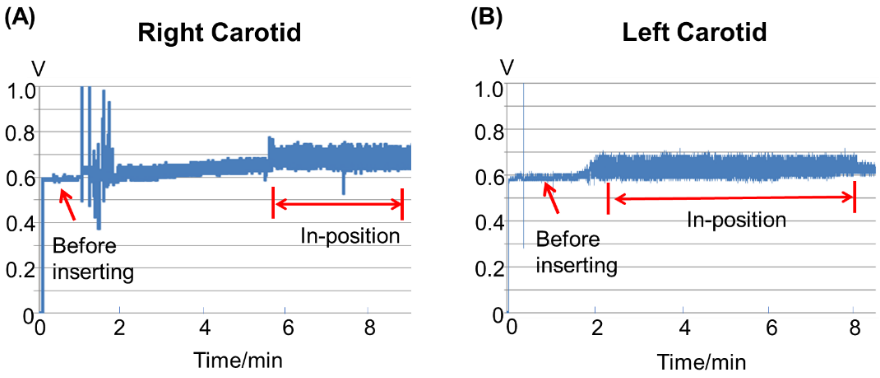


**Fig. SI-1: FFR pressure measurements**. Pressure changes were recorded from both the right and left carotid arteries upon catheter deployment to the target vessel.

*SI-2. Comparison of impedance measurement with the computational model*

From each carotid artery sample, we created a 3-D model and scanned through possible electrode positions with a different combination of $z$ and $\theta$. 15-permutation impedance values were calculated for each combination, and computational EIS profiles were compared with the measured EIS to identify the best fit scenarios. Scatter plots of the 15-permutation impedance from the measured EIS and representative computation model (simulation) are presented with the best fit combination of $z$ and $\theta$ as highlighted in light blue (**Fig. SI-2**).

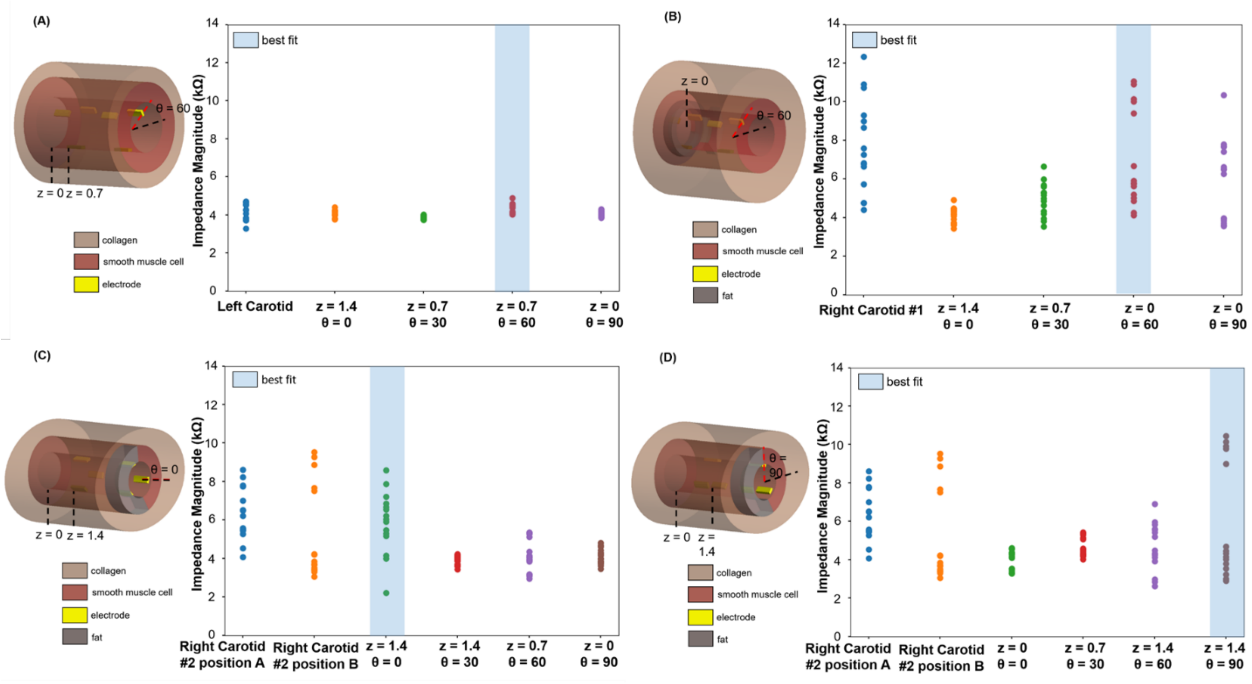


**Fig. SI-2: Computation Models**. Comparisons with the simplified 3-D schematic indicate the position of the electrodes in the endolumen where the computational impedance values are overlapping with the experimental EIS. Experimental impedance at 10 kHz is plotted alongside the modeling results at the given z and θ values, and the combination with the best fit was highlighted. Computational EIS are compared among (A) left carotid artery, (B) RC1, (C) RC2 position A, and (D) RC2 position B.

### *SI-3. Custom-built circuit for interfacing with commercial FFR probe*

The pressure measurement was performed via a Wheatstone bridge circuit (**Figure SI-3**). Changes in pressure affect the simultaneous increase or decrease of the two resistors (R±ΔR) that attached onto the membrane structure of the sensor. By maintaining the equality between R1 and R2 ($R_1 = R_2$), the reading of $V_{out}$ was linearly dependent on the change in resistance (ΔR) (*56*).

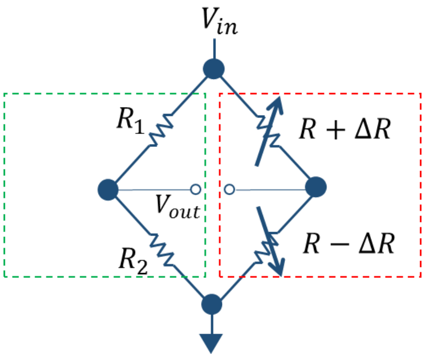


**Fig. SI-3**: **Schematics of the circuitry used in FFR adaptation**. The half bridge inside the red dotted square represents the components integrated in the commercial FFR probe, and the half bridge inside the green dotted square represents the standard resistors.

*SI-4. Comparison of impedance measurement before and after electroplating*

The flexible electrodes (either before or after electroplating) were submerged in a large container of saline solution (0.9% wt NaCl). The geometric effect was considered negligible under these settings. The impedance spectra were measured using Gamry G 300 with 50 mV, and the frequency ranged from 100 Hz to 300 kHz. The electroplating of Pt Black resulted in a low contact impedance for the electrodes, as evidenced by the significantly flattening in the EIS curve beyond 1 kHz. Therefore, we validated the selection of 10 kHz for the EIS analyses and computational modeling.

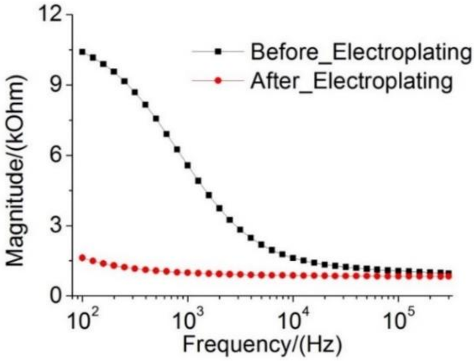


**Fig. SI-4**: **Electrochemical impedance spectra.** A comparison between before and after electroplating demonstrated a significant decrease in contact impedance.

*SI-5. Reconstruction of conductivity mapping from the impedance data*

Our computational model was composed of the arterial wall that was represented by 576 elements (**Figure SI-5Bi**), and the annular collagen layer by 288 elements (**Figure SI-5Bii**). The initial conductivities of arterial wall elements were derived from the EIS measurements. The collagen layer was assigned with a large element size and a uniform conductivity. We varied the conductivity distribution to a direction that optimally reproduced the 15 permutations for the impedance measurements. To this end, we used the "genetic algorithm" so that all elements "evolved" to reach the final mapping results. The implementation was as follows: the conductivity value from each of the 864 elements was considered as an 864×1 vector. We generated a solution candidate pool (100) by adding a Gaussian-distributed noise to this initial 864×1 vector, $\vec{C}_1, \vec{C}_2, \dots \vec{C}_{100}$. For each candidate, we calculated the 15 impedance values by solving the Laplace equation:

$$Z_{12,\mathrm{sim}}, Z_{13,\mathrm{sim}}, Z_{14,\mathrm{sim}}, \dots Z_{56,\mathrm{sim}} \quad (1)$$

We defined our fitness function as follows:

$$f = \sqrt{(Z_{12,\mathrm{measured}} - Z_{12,\mathrm{sim}})^2 + (Z_{13,\mathrm{measured}} - Z_{13,\mathrm{sim}})^2 + \cdots (Z_{56,\mathrm{measured}} - Z_{56,\mathrm{sim}})^2} \quad (2)$$

The "genetic algorithm" was implemented by the following steps:

i. Calculate the fitness function for all of the solution candidates in the pool.

ii. Rank the candidates according to their fitness function from small to large values.

iii. Identify the top-10 candidates from the pool: $\vec{C}_1^*, \vec{C}_2^*, \dots \vec{C}_{10}^*$.

iv. Generate a new pool of 90 candidates as follows:

$$\vec{C}_i^* = \sum_{k=1}^{10} (\sigma \lambda_k \vec{C}_k^* + (1-\sigma)\, \lambda_k \frac{\sum_1^{10} \vec{C}_k^*}{10}), i = 11,12, \dots 100, \quad (3)$$

where $\sigma = 1.3$ is a factor to moderate the boundaries of the candidate space obtained from the above equation, and $\sum_{1}^{10} \lambda_k = 1$ (*50*).

Steps i-iv were repeated until the minimum fitness function reaches the predefined target, and the solution candidate $\vec{C}_1^{final}$ was used to assign to the conductivity distribution to generate the final EIT mapping (see **Figure 3**). We excluded the outmost layer as the calculated conductivity distribution was fairly homogeneous since the major heterogeneity resided in the inner layer (576 elements).

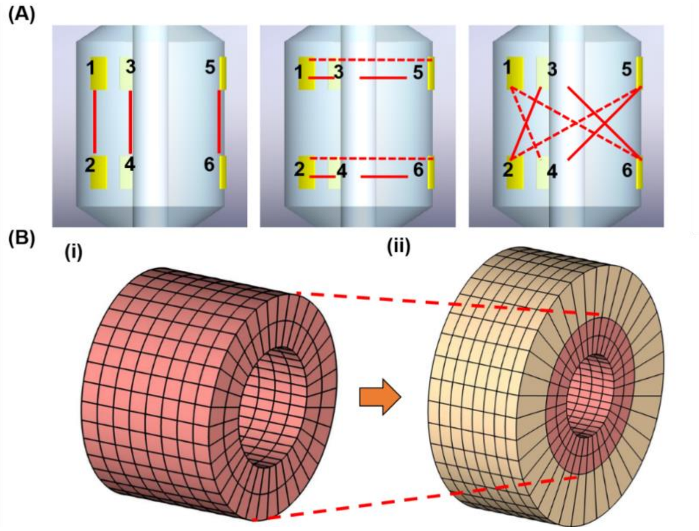


**Fig. SI-5: Finite element model for reconstructing the conductivity maps.** (A) The configuration of the 6-point electrodes (EIS sensor) generates 15 permutations. (B-i) 576-element mapping scheme represents the inner smooth muscle layer. (B-ii) 864-element mapping scheme includes the collagen layer (yellow).

*SI-6. Histology-based 3-D FEM modeling*

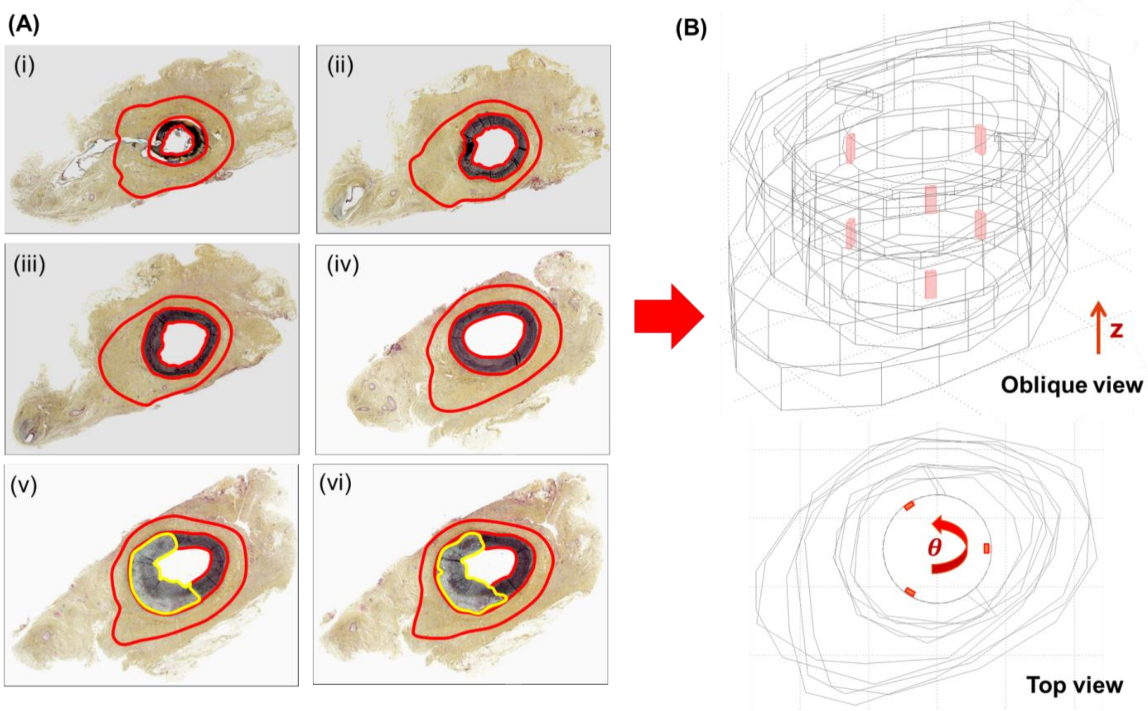


**Fig. SI-6: Finite Element Model for 3-D histology and computational modeling.** (A) Representative histological cross-sections from the right carotid artery (RC2) were demarcated by the Movat staining for connective tissue, the boundaries for collagen, smooth muscle, and the lipid component. (B) These demarcations from the histological slices allowed for reconstructing a 3-D model. The positions of the electrodes (red) were defined by the $z$ and $\theta$ coordinates.

**Table I. Tissue properties used for the computational model (57)**

| | Collagen layer | Smooth muscle cell layer | Fat tissue |
|---|---|---|---|
| $\boldsymbol{\sigma}$ $(\boldsymbol{S/m})$ | 0.174 | 0.307 | 0.042 |
| $\boldsymbol{\varepsilon}$ | 32000 | 149000 | 193000 |
